# An Arabidopsis leaf expression atlas across diurnal and developmental scales

**DOI:** 10.1101/2023.11.10.566572

**Authors:** Gina Vong, Kayla McCarthy, Will Claydon, Seth J. Davis, Ethan J. Redmond, Daphne Ezer

## Abstract

Mature plant leaves are a composite of distinct cell types, including epidermal, mesophyll and vascular cells. Notably the proportion of these cells, and the relative transcript concentrations within different cell types may change over time. While gene expression data at a single-cell level can provide cell-type specific expression values, it is often too expensive to perform this on high resolution time series. Although bulk RNA-seq can be performed in a high resolution time series, the RNA-seq in whole leaves measures the average gene expression values across all cell types in each sample. In this study, we combined single cell RNA-seq data with time-series data from whole leaves to infer an atlas of cell type-specific gene expression changes over time for *Arabidopsis thaliana*. We inferred how relative transcript concentrations of cell types vary across diurnal and developmental time scales. Importantly this analysis revealed three sub-groups of mesophyll cells that have distinct temporal profiles of expression. Finally, we develop tissue-specific gene networks that form a new community resource: An Arabidopsis Leaf Time-Dependent Atlas (AraLeTa), which allows users to extract gene networks that are confirmed by transcription factor binding data and specific to certain cell types, at certain times of day and certain developmental stages, which is available at: https://regulatorynet.shinyapps.io/araleta/.

## Introduction

The coordination of spatial and temporal gene expression dynamics is fundamental to plant development and response to environmental stimuli. Organisms have distinct gene-regulatory programmes within different cell types, which regulate the changes in gene expression over diurnal (Yakir et al., 2011) and developmental time scales (Ma et al., 2005). Each of these regulatory programmes coordinate changes in gene expression over time to respond to both intrinsic (time of day, maturity) and extrinsic factors (environmental stimuli) (Sheen, 1994). It is the coordination of transcriptional patterns in discrete cell types that provides determinate capacity for cell function in a context of its tissue and organ.

Multiple approaches have been taken to measure gene expression over space and time, but each have their drawbacks. Through live image-based assays, it is possible to track gene expression of a small number of genes over both time and space (Shav-Tal et al., 2004). For instance, Gould et al. identified waves of circadian gene expression originating from the meristems (2018). In contrast, RNA-seq enables researchers to measure gene expression of all mRNAs at once. Many researchers perform high temporal resolution RNA-seq time series experiments to infer how gene expression changes over time (*e.g.* Balcerowicz et al., 2021, Krouk et al., 2010, Cortijo et al., 2017, Ezer and Keir, 2019). However, these studies do not capture the cell-type specific changes in gene expression. Additionally, over 80% of leaf expression in a bulk RNA-seq sample originates from mesophyll cells, which will mask gene expression patterns from other cell types, such as in the vasculature and epidermis tissues (Endo et al., 2014). Bulk RNA-seq also masks heterogeneity within mesophyll cell populations (Procko et al., 2022). Increasingly, single cell and tissue specific RNA-seq has been used to characterise cell-type specific gene expression patterns in leaf (Kim et al., 2021; Lopez-Anido et al., 2021; Liu et al., 2020), root (Shahan et al., 2022) and meristem (Neumann et al., 2022) tissue, but it is prohibitively expensive to perform these in a high-resolution time series. Lee et al. (2023) has developed a developmental single cell atlas, but it only covers five vegetative stages and is therefore not at a similar temporal resolution as existing bulk RNA-seq resources and would be too expensive to replicate under a wide range of experimental conditions. Recent efforts are underway to construct further Plant Cell Atlases based on single cell analysis (Ahmed et al., 2021), and it is important to find methods to best utilise these kinds of resources, especially when investigating processes that occur over time.

Our work here demonstrates the potential of integrating single-cell RNA-seq data and high-resolution time series data to unravel cell-type specific gene networks in *Arabidopsis thaliana*. Drawing inspiration from cancer cell dynamics research (Newman et al., 2019), we used CIBERSORTx, a powerful technique that combines single-cell RNA-seq and bulk RNA-seq data (Newman et al., 2019) that outperforms other deconvolution methods (Sutton et al., 2022). We inferred relative expression of various cell types across samples and estimated cell type-specific gene expression values. By applying these techniques, we gained insights into the dynamics of expression within different cell types across varying temporal scales. Although Procko *et al*. (2022) identified four sub-populations of mesophyll with unclear distinguishing markers, our analysis reveals changes in their relative expression levels over diurnal and developmental scales, raising questions about cell-state changes and activity-level variations over time, as well as the relative light sensitivity of different mesophyll cell states. Moreover, differences in relative expression of mesophyll subgroups between bolted and unbolted plants were observed. Through incorporating transcription factor binding data (O’Malley *et al*, 2016), we have provided a valuable resource for the Arabidopsis community: leaf-cell type network available at https://regulatorynet.shinyapps.io/araleta/. We also highlight relevant portions of this network during diurnal and developmental time scales.

## Results

### Detection of cell type transcriptional activity in bulk RNA-seq by utilising single cell RNA-seq data

First, we wished to confirm that we can accurately predict proportions of cell types in bulk RNA-seq samples using single cell RNA-seq data from *Arabidopsis thaliana* (**Fig 1A**), by training CIBERSORTx on a training set of single cell RNA-seq cells and then testing its accuracy on deconvolving simulated bulk RNA-seq samples constructed from subsets of the remaining cells. When we simulated bulk RNA samples containing a single cell type, we correctly identified all tissue types, except for hydathodes, as these were often mis-classified as mesophyll cells (**Fig S1**). In this paper, we named the three mesophyll cell clusters (specifically, clusters 1, 3, and 4) from Procko *et al*. (2022) as mesophyll groups 1, 2 and 3, respectively. We also named the three clusters of unknown type (termed clusters 10, 11, and 16) as unknown groups 1, 2 and 3, respectively. Next, we simulated bulk RNA-seq samples with mixed cell types. We accurately predicted the relative abundance of simulated cell types between these samples (**Table S1**), with Pearson’s R greater than 0.8 for all cell types except for two unidentified cell types and sieve cells (**Fig S2, Fig 1B**). However, we note that CIBERSORTx is best at predicting the relative amounts of cell types between samples, rather than the relative proportions of cell types within a sample – a known issue with gene expression deconvolution algorithms (Sutton el al., 2022). It consistently over-estimated the proportion of stressed cells and underestimated the proportion of other cell types (**Fig 1C**). These results confirm that CIBERSORTx, a technique initially developed for mammalian research (Newman et al., 2019), can be applied to plant systems.

**Figure 1:**
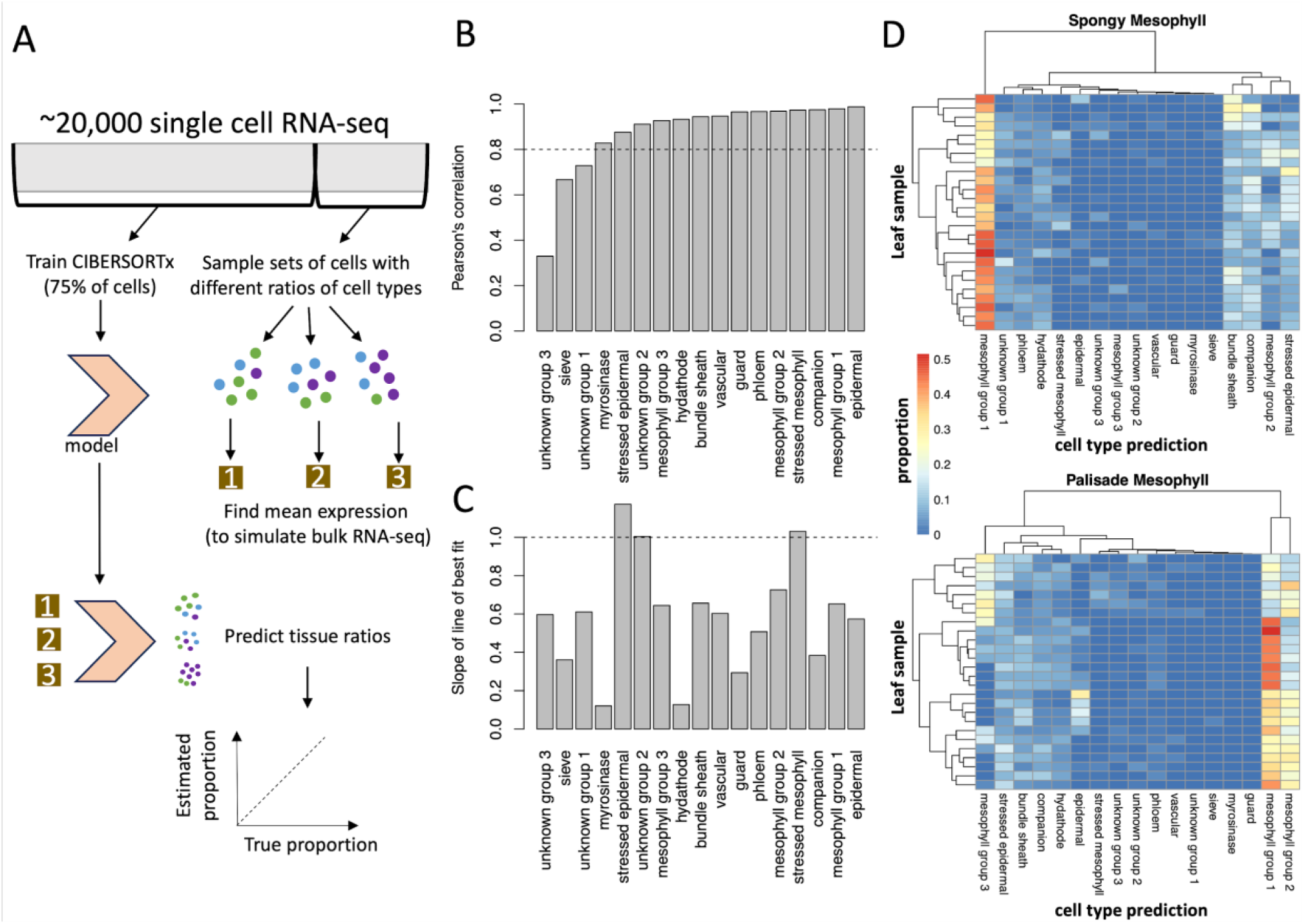
CIBERSORTx predictions for simulated bulk RNA-seq samples. (A) 75% of the cells per cluster were used to train a CIBERSORTx model, while the remaining 25% of cells were repeatedly sub-sampled to generate 500 simulated bulk RNA-seq samples with mixed cell types (see Methods). (B) The Pearson’s correlation between the true cell type proportions in the simulated bulk RNA-seq sample and the tissue proportions predicted by CIBERSORTx. All the correlations were statistically significant (p<0.05), with Pearson’s correlations >0.8 highlighted by the horizontal bar. (C) The slope of the line of best fit for the comparison between the predicted and true cell type proportions (scatterplots in Fig S2). Values close to 1 (horizontal line) are most accurate. Taken together, these results suggest that CIBERSORTx can predict the relative proportions within the same cell type, between RNA-seq samples, but that it consistently over- or under-predicts certain cell types within an RNA-seq sample. (D) Predicted cell type proportions in microdissected leaf samples, with each row representing a leaf and each column representing a different predicted cell type. Note that phloem is short for phloem parenchyma.

CIBERSORTx was also successful at predicting the tissue composition of simulated bulk RNA-seq samples generated from other single cell RNA-seq experiments (**Fig 1D, Fig S3**). Interestingly, microdissected spongy mesophyll cells from Xia et al., (2022) tended to be classified as mesophyll cells from group 1, while the palisade cells were more evenly split between the three mesophyll groups that Procko et al. (2022) previously reported (**Fig 1D**). This suggested that atlases of single cell RNA-seq expression can serve as reusable resources in the community for estimating cell type compositions of bulk samples.

Next, we explored the parameters of the signature matrix constructed from the training set to help with deconvolving the bulk RNA-seq samples. CIBERSORTx selected 4950 genes for use in predicting the cell-type proportion (**Fig S4**), and these have significant enrichment for Gene Ontology (GO) terms associated with ion binding, catalytic activity, structural constituents of chromatin, transmembrane transporter activity, response to stimulus/stress and amino acid metabolic processes-- See **Table S2 (Signature matrix), S3 (gProfiler outputs).** These cell-type specific processes appear to be capable of distinguishing cell type proportions.

### Across developmental scales, transcriptional activity shifts from epidermal to vascular cell types

We next sought to identify how cell-type transcriptional activities changed over different temporal scales. For this we applied the signature matrices that we previously inferred from single cell RNA-seq data to interpret bulk RNA-seq time series datasets. It is important to note that deconvolution algorithms like CIBERSORTx do not find the proportion of cells of each type, but rather the proportion of transcripts that is attributable to different cell types. Changes in cell-type proportion over time could be attributable to changes in the total number of cells at a certain time, increased activity of certain cell types, or cells changing their expression profile to mimic the expression of a different cell type. For conciseness, we will refer to the output of CIBERSORTx as the “activity” of specific cell types.

Next, we investigated whether the proportion of cell type expression levels varied across a developmental time scale (**Fig 2A, Fig S5, Table S4**), utilising a leaf developmental time series RNA-seq dataset (Woo et al., 2016). The reference single cell RNA-seq experiment was performed on the first true leaves at 17 days post-germination, so it only captures cell type specific transcriptional activity at a single snapshot. During the growth-to-senescence transition, we detected a decrease in epidermal expression and an increase of vascular expression. Rarer cell types, like guard cells and hydathodes, seemed to have higher proportion early in development, possibly because these cell types get diluted as the leaf expands (Ietswaart et al., 2017) or because these rarer cell types have gene expression profiles that mimic other cell types when mature, as previously shown for guard cells (Adrian et al., 2015). Phloem parenchyma expression and stressed mesophyll expression was maximised in the senescing leaves compared to other developmental stages. This is consistent with phloem parenchyma cells contributing to nutrient redistribution in the senescing leaf and with the transformation of neighbouring cell types into phloem cells during senescence (Hunziker et al., 2019).

**Figure 2:**
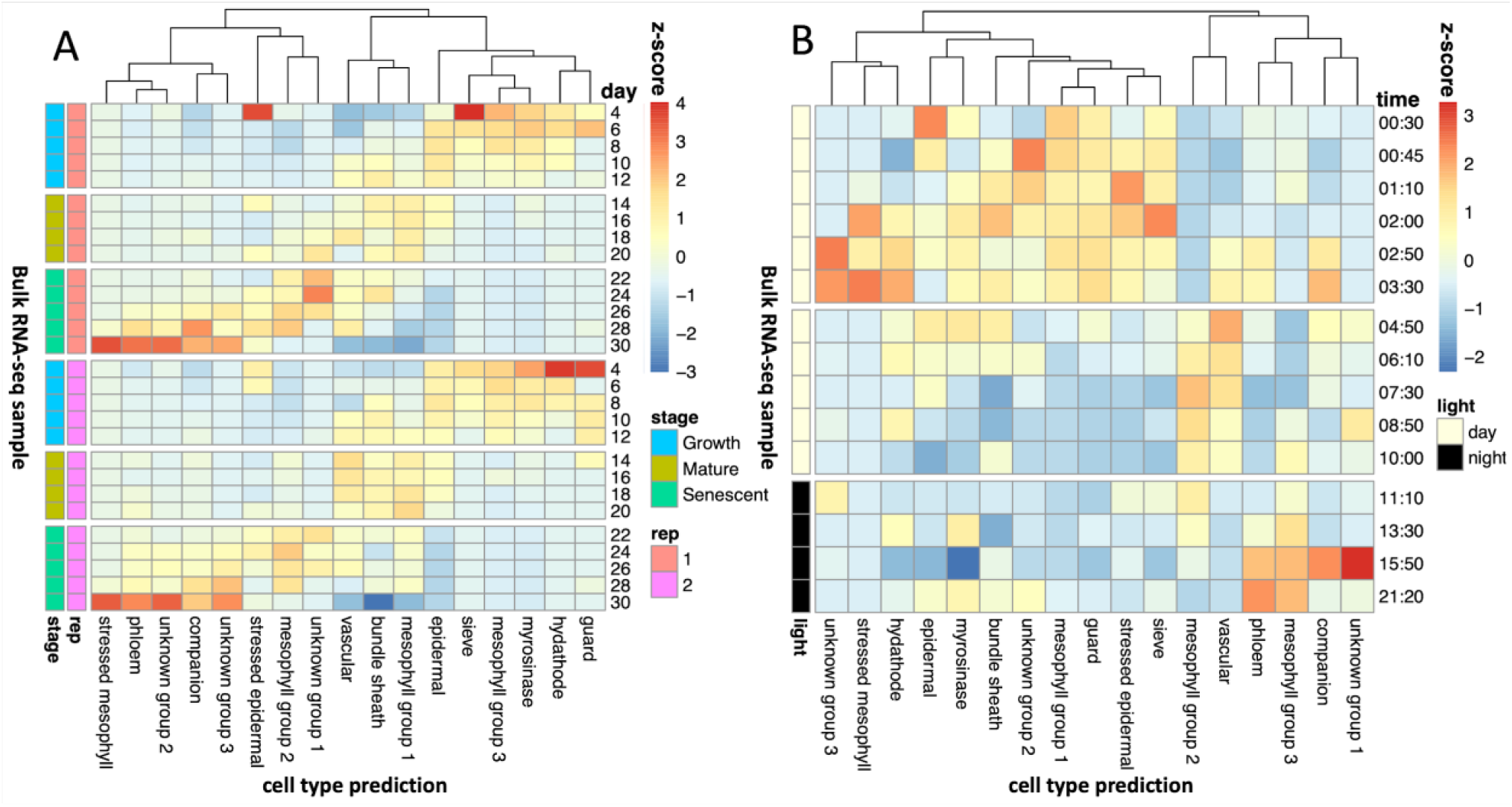
Shifts in cell type proportions over developmental and diurnal time series. We predicted the proportion of cell types in a (A) developmental time series (Woo et al., 2016) and (B) diurnal time series (Hickman et al., 2017), utilising a reference leaf scRNA-seq dataset (Procko et al., 2022). In (A), each row represents a different RNA-seq sample, with the proportions normalised using z-scores. The developmental stage and the biological replicate are indicated by the coloured bars. In (B), the proportions of cell types were predicted independently for each of the 4 biological replicates, and then these were averaged for each row, then normalised by z-scores. The times are relative to dawn.

Intriguingly, the three groups of non-stressed mesophyll cells also displayed distinct temporal patterns. Group3 mesophylls have peak expression during the growth phase, group 1 have peak expression at the mature leaf phase and group 2 have peak expression during the mature-leaf-to-senescence transition period. Group 1 cells were found most frequently in the leaves in Procko et al. (2022), which is consistent with their sample collection time point of 17 days. This analysis suggests that mesophyll-specific expression profiles transition between the three different states identified by Procko et al., (2022) over developmental time scales.

### Diurnal oscillations of mesophyll, epidermal, vascular, and phloem transcriptional activity

Additionally, we were interested in how the cell-type specific gene expression patterns varied across diurnal time scales, as measured in a leaf diurnal time series RNA-seq data set (Hickman et al., 2017), see **Fig 2B, Table S5**. Procko et al. (2022) grew their leaves under continuous light to minimise the impacts of the clock, but plant clocks are synchronised by the initial seed imbibement and so they will still experience consistent daily oscillations (Zhong et al., 1998). Even if averages of transcripts appear “arrhythmic” after prolonged plant acclimation to constant conditions, the cells that pattern tissues are still robustly rhythmic, albeit asynchronous from each other (Yakir et al., 2011; Gould et al., 02018). Thus, the individual cells sampled by Procko et al, (2022) will each be in an unknown phase of the circadian clock.

Epidermal expression is maximised in the morning. Meanwhile, vascular expression peaks in the afternoon. We hypothesise that this is when the plant is most water-stressed, due to the tendency for this time of day to have a higher temperature. Phloem parenchyma expression peaks at the end of the night when passive loading of sugars through plasmodesmata is replaced with active apoplasmic loading (Wei et al., 2021). Interestingly, epidermal stress cell expression peaks during the ZT1 dawn burst of expression (Balcerowicz et al., 2021), while mesophyll stress expression peaks a few hours afterwards (ZT2-4). One hypothesis is that epidermal cells become stressed by the lights suddenly turning on within the growth cabinet, while mesophyll cells become stressed because of ROS accumulation as result of photosynthesis.

The three non-stressed mesophyll groups primarily have peak expression at different times of day. The mesophyll cells that are predicted to be active during early development (group 3) are also predicted to be active late at night. The mesophyll cells that were active in mature leaves (group 1) were also active in the morning. A comparison with Xia et al. (2022) suggests that this mesophyll group may be enriched for spongy mesophyll cells. The mesophyll group that was most active during the transition to senescence (group 2) is also active during the afternoon and early evening. The mesophyll sub-groups each appear to have distinct temporal transcriptional profiles, both on a diurnal scale and developmental scale.

### Expression of time-of-day dependent light sensitive genes across cell types

Next, we evaluated whether the cell types had different levels of expression of light responsive genes. We hypothesised that group 1 mesophyll cells may include more of the genes that are induced by light exposure at the end of the night, as these cells are most active in the morning. To evaluate this hypothesis, we analysed the expression of light-induced genes (Rugnone et al., 2013) in the Procko et al., (2022) single cell data set. Consistent with our hypothesis, genes that are induced by light exposure at night (Fig 3A) tended to be found in mesophyll group 1 (morning/mature phase mesophyll group). This set of mesophyll group 1 genes included NIGHT LIGHT–INDUCIBLE AND CLOCK-REGULATED1,3 (LNK1, LNK3), which help entrain the circadian clock in response to light in the morning (Xie et al., 2014), SALT TOLERANCE (STO), which is a BBX family protein involved in the morning dawn burst of expression (Balcerowicz et al., 2021) and members of the light harvesting complex in photosynthesis (LHCB2.4, LHCA4). Thus mesophyll clusters indeed harboured notable light-induced genes.

**Figure 3:**
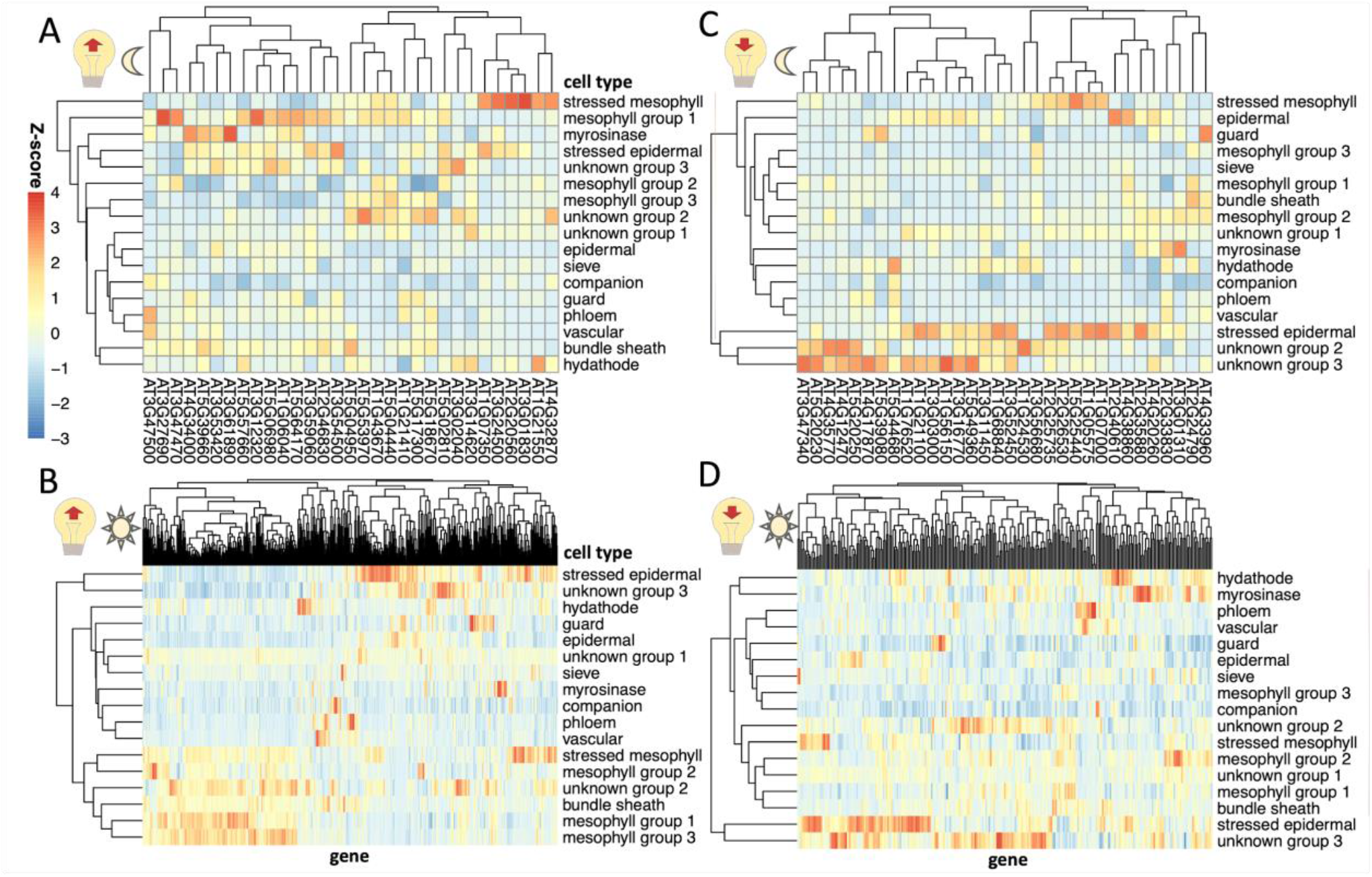
Expression of light sensitive genes in single cell leaf data. Sets of genes were selected that are light-induced (A, B) or light-repressed (C, D) during either a nocturnal light treatment (A, C) or light treatment after an extended night (B, D), based on Rugnone et al. (2013). The mean expression of these genes in cells in each cluster in Procko et al. (2022) were calculated and a z-scores was calculated for each column.

A different set of genes that are light sensitive at night were more highly expressed in stressed mesophyll cells. These genes include DNAJ, a heat shock protein (Pulido and Leister, 2017), SERINE/ARGININE RICH-LIKE PROTEIN 45A (SR45a), a stress-induced splicing factor that regulates anthocyanin accumulation (Gulledge et al., 2012; Albaqami, 2023), and MULTIPROTEIN BRIDGING FACTOR 1C (MBF1C) whose expression is elevated in response to a wide range of stresses (Lee and Bailey-Serres, 2019). These results suggest that light exposure earlier than anticipated may induce a light stress response, in addition to activating morning mesophyll expression, and that these two responses impact the activity of two different sets of genes in two different cell types. In contrast, genes that are induced by light exposure after an extended night (Fig 3B) are primarily either expressed in the mesophyll or are expressed in stressed epidermal cells.

Despite the plants in Procko et al. (2022) being exposed to continuous light, some cell types continue to express genes that are repressed under light exposure. Specifically, unknown groups 2 and 3 contain genes that have decreased expression under light exposure at night like SENESCENCE1 (SEN1), DARK INDUCIBLE 10 (DIN10) and GLUTAMINE-DEPENDENT ASPARAGINE SYNTHASE 1 (ASN1), which are all also induced by senescence (Fujiki et al., 2001). Coupled with our previous observations that their activity peaks during late senescence (Fig 2A), we propose that these cell types are associated with senescence. Stressed epidermal cells also contain high levels of light repressed genes. Genes that have reduced expression under light exposure are not expressed highly in the cell types that are predominantly found enriched during the night (phloem, mesophyll group 3, and companion cells, Fig 2B), but this may be because the scRNA-seq was performed in plants under continuous light (Fig 3C,D). This suggests that it may be wise to perform scRNA-seq on more realistic diurnal conditions, sampling multiple times a day, to adequately capture the cell specific transcription over the course of a day.

### Cell type-specific regulatory programme during bolting

We have observed that there are differences in the transcriptional activity of cell types across a developmental and a diurnal time scale. Next, we decided to focus on the changes that happen during a rapid developmental transition, specifically bolting, which coincides with the start of senescence (Redmond et al., 2023). Mimicking the pattern observed during from the developmental time series Fig 2A, the relative expression of mesophyll group 3 (the growth-phase mesophyll) goes down, while the expression of group 1 and 2 mesophyll (the mature-phase and senescence mesophyll) went up over pseudotime (Fig 4A, Table S6), where pseudotime refers to the predicted ordering of individual plants in Redmond et al. (2023) over a developmental trajectory on the basis of bulk RNA-seq data (Fig 4B, Table S6). Also consistent with the developmental time series, we observe a decrease in epidermal and sieve cell activity. The consistency between these cell type activity predictions between the developmental atlas (Woo et al., 2016) and the bolting pseudotime (Redmond et al., 2023) suggests that these developmental shifts in cell activity are robust and confirms that the pseudotime is effectively ordering by a developmental trajectory.

**Figure 4:**
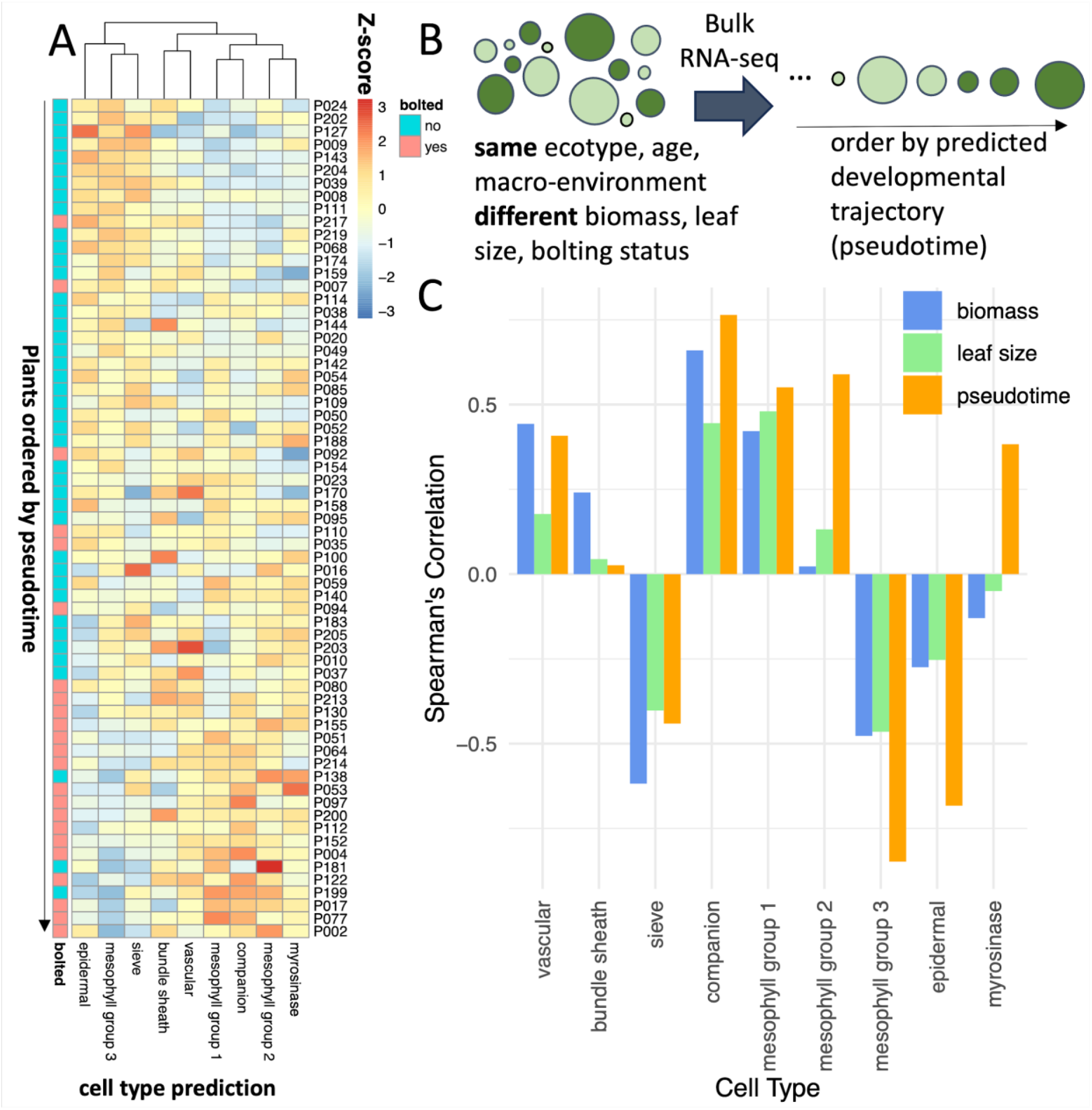
Cell activity changes during bolting. (A) We predicted the proportion of cell types in plants immediately before and after bolting (Redmond et al., 2023), utilising a reference leaf scRNA-seq dataset (Procko et al., 2022). Each row represents a different RNA-seq sample (representing a single plant), with the proportions normalised using z-scores. (B) The plants in (A) are ordered by pseudotime, which was calculated by Redmond et al., (2023) as an arrangement of the 65 individual plants sampled along a developmental trajectory. (C) For each cell type, the Spearman ranked correlation was calculated between the proportion of that cell type in each plant vs another trait of the plant (biomass, leaf size, or pseudotime).

We were curious whether the changes in cell types over time were associated more with the plant’s development or with its other traits, such as biomass or leaf size. For each cell type, the Spearman’s correlation coefficient was calculated between the proportion of that cell type in each plant and that plant’s pseudotime, wet biomass or leaf area (Fig 4C). The proportion of vascular cells, bundle sheath, and sieve cells were more strongly correlated with biomass than pseudotime, suggesting that the size of the plant may be associated with water and sugar transport. On the other hand, mesophyll, epidermal and companion cells are more associated with the pseudotime and so may be more closely associated with development.

Next, we investigated the cell type-specific processes taking place during bolting, utilising the capacity of CIBERSORTx to impute cell type specific expression from bulk RNA-seq data. The cell type specific genes increased their expression at different points in the pseudotime using the bulk RNA-seq data from Redmond et al., (2023) (Fig 5A), suggesting that at least part of the differences in the timings of expression of genes across pseudotime could be a result of different activity levels of different cell types. Most of the genes were identified as being associated with mesophyll group 2 (Fig 5B). There are some consistencies with the mean expression levels of these genes in the scRNA-seq data set (Procko et al., 2022), specifically amongst mesophyll group 2 cells and companion cells for genes that increase with pseudotime and with mesophyll group 2/3 and sieve cells for genes that decrease with pseudotime (Fig 5C). However, there are also some inconsistencies, such as being unable to reconstitute the epidermal expression pattern. Over all genes, the imputed expressions are relatively consistent with the known gene expression values (Fig S6).

**Figure 5:**
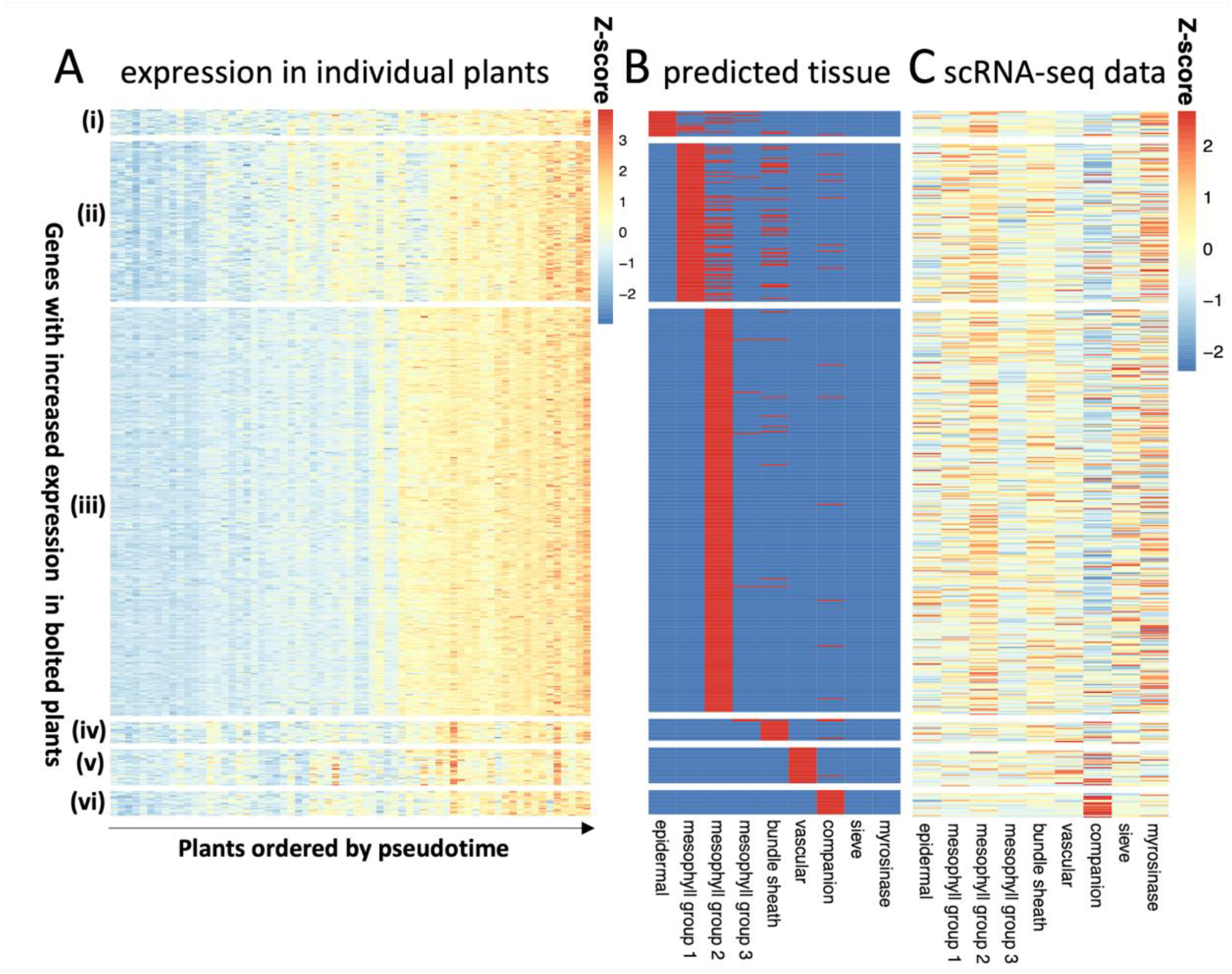
Cell-type specific gene expression of bolting-related genes. Using the high-resolution imputing function in CIBERSORTx, we predicted cell-type specific expression of genes that were differentially expressed in bolted/unbolted plants. Here we show the results for the genes that have higher expression in bolted plants, with the inverse gene set shown in Figure S8. (A) shows the z-score of the expression of these genes in the bulk RNA-seq experiment (Redmond et al., 2023), grouped by the cell type in which they were predicted to be expressed and ordered by pseudotime. (B) shows the cell type assignment, with red indicating that a gene is expected to be found in that cell type. (C) shows the z-score of the mean expression of these genes in the scRNA-seq data set (Procko et al., 2022).

First, we analysed the GO terms of cell type-specific genes whose expression increased over pseudotime (**Table S7**). Those associated with mesophyll group 2 tended to be found in membranes and involved in ATP, GTP and NAD+ binding (GO:0005524, GO:0005096, GO:0003953) and regulation of endocytosis (KEGG:04144). Genes found in the epidermis were enriched vesicle-mediated transport (GO:0016192). Vascular genes were associated with sulphur metabolism (KEGG:00920) and response to nutrient levels (GO:0031669).

Among genes that reduce their expression over pseudotime (**Table S8**), there was enrichment in chloroplast and photosynthesis-related GO terms in epidermal and mesophyll cell types. Epidermal cells also were enriched in porphyrin (a pigment) metabolism (KEGG:00860) and amino acid metabolism-related processes (KEGG:00300). Mesophyll group 3 was associated with the cell cycle (GO:0007049) and DNA replication initiation (GO:0006270), while both groups 2 and 3 were enriched for components of the ribosome (GO:0005840) compared to other cell types. We noted that mesophyll group 3 is also expressed in early development and at night, so potentially this cell type is in a dividing and growing state.

### AraLeTA: An Arabidopsis Leaf Time-dependent Atlas

Our previous results have demonstrated that there are different regulatory programmes that are active at different times of day, different developmental stages, and different cell types. As a resource for the community, we have developed AraLeTA, and Arabidopsis Leaf Time-dependent Atlas (https://regulatorynet.shinyapps.io/araleta/), which can be used to identify the portions of the Arabidopsis gene regulatory network that are active in different contexts, by filtering based on expression values (Fig 6). As its basis, it utilises the DAP-seq network developed by O’Malley et al., (2016), but it enables the user to filter the network by the age, time of day, and cell types of interest, highlighting edges in which both the source and target are expressed above a threshold value under the relevant conditions. This kind of thresholding approach uses similar criteria for filtering the gene regulatory network as what was used by Ferrari et al., (2022) and has been shown to enrich the gene network for true edges. The AraLeTA network can be downloaded or visualised as a heatmap or graph.

**Figure 6:**
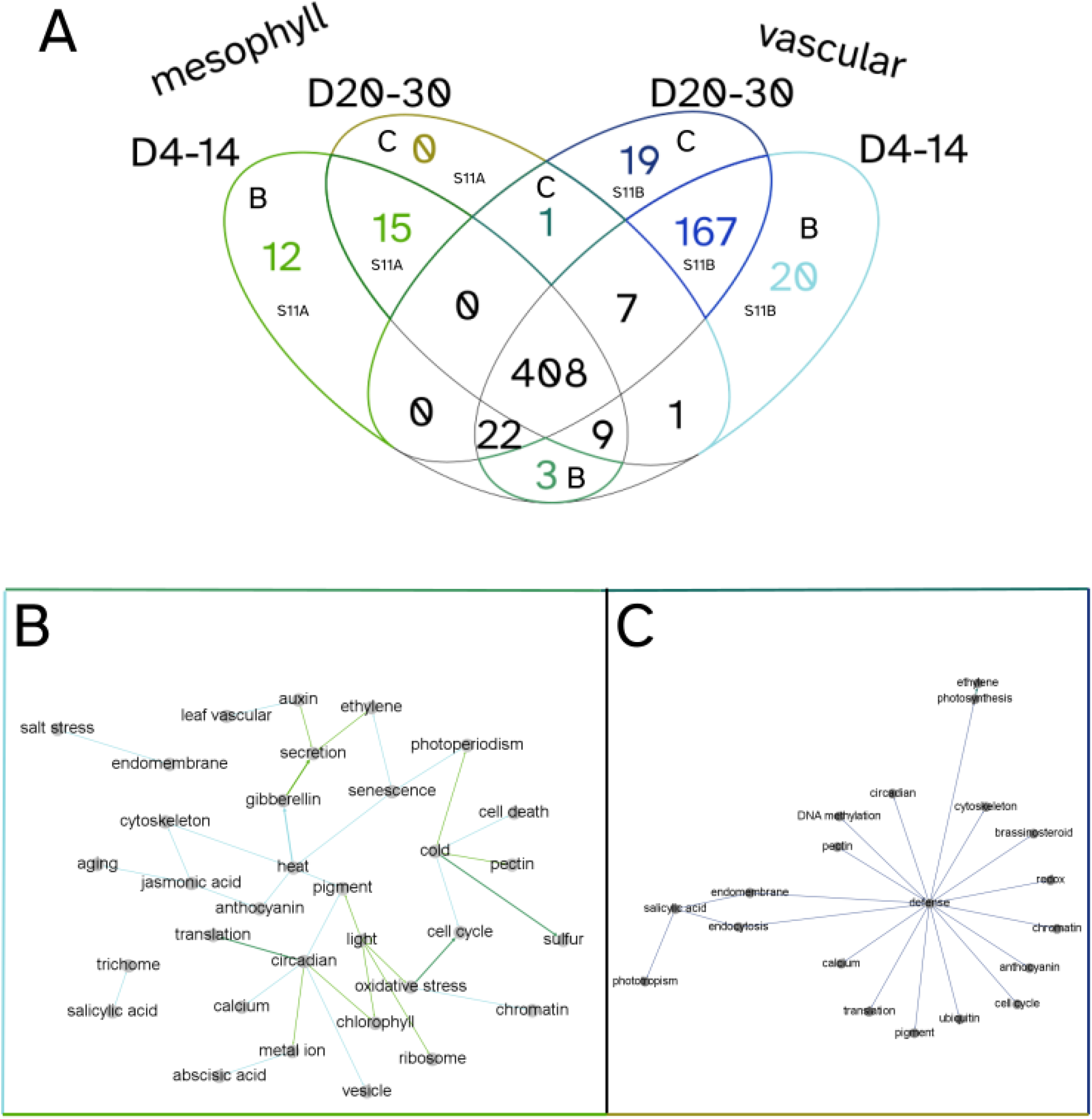
PAFway networks across different tissue types at different ages. Using PAFway, we generated networks of functional terms to assess changes in function between tissue-types as they age. (A) Mesophyll (groups 1, 2, 3) and vascular associated (bundle sheath, phloem, vascular, companion, guard and sieve) cells showed a large overlap in functional edges across age-groups, with some network edges which are unique to specific tissues and ages. (B) Network of functional terms which are associated with young mesophyll and vascular cells. (C) Network of functional terms associated with old mesophyll and vascular cells. Together, these show the changing network over time, suggesting that these cells perform different physiological functions as they age. The colours of the edges in (B) and (C) correspond to the associated coloured sections of the Venn diagram in (A).

As it is often difficult to make sense of large “hairball” networks, we have incorporated PAFway additionally, a programme for identifying the relationships between functional terms within the topology of the network (Mahjoub and Ezer, 2020). This allows us to find patterns in the rewiring of the network over time, within specific cell types.

To highlight the utility of AraLeTA, we focussed on four conditions: (i) young (4-14 days post germination) mesophyll (groups 1, 2, 3) cells (ii) old (20-30 days post germination) mesophyll cells (iii) young vascular (bundle sheath, phloem, vascular, companion, guard and sieve) cells and (iv) old vascular cells (Fig 6A). To select thresholds for our networks, we varied the single cell and bulk RNA-seq threshold parameters and evaluated their impact on the number of significant functional edges selected by the PAFway network, choosing thresholds that generated either local maxima or inflection points in the size of the PAFway network (Fig S10). As expected, young mesophyll cells included many associations with light response (Fig 6B). Interestingly, there are more defence, jasmonic acid (JA) and salicylic acid (SA) related edges in vascular cells, reflecting the transport mechanism of these plant hormones (Fig S11). The old plant networks are centred on SA, a senescence associated plant hormone, and defence (Fig 6C). Many of the genes with GO terms related to defence are also involved in senescence, especially in response to necrotrophic pathogens (Woo et al., 2016). Redmond et al. (2023) also demonstrated that processes such as ubiquitination, endocytosis, cell cycle, translation and response to redox are all perturbed at the onset of senescence, and these terms all appear in the network associated with older plants. These general trends confirm that the filtering criteria are adequately selecting relevant portions of the DAP-seq network.

## Discussion

### Cell type specific transcriptional activity: what does it mean?

Here we illustrated how we used CIBERSORTx to infer the transcriptional activity of different cell types within a bulk RNA-seq sample. One must be nuanced in our interpretation of what a change in transcriptional activity means from a biological standpoint (Fig S12). When one cell type is predicted to have a higher proportion than another cell type, there are many alternative explanations: (i) There may be a higher proportion of one cell type in the sample relative to the other, (ii) One cell type may have a higher overall transcription rate than the other or (iii) There may be only one cell type that fluctuates between transcriptional states. For instance, it is unclear whether it is possible for cells that belong to a particular mesophyll group to transition into a cell from a different mesophyll group; (iv) cells may express a transcriptional pattern that is similar to other cell types at certain time points and the algorithm may be misassigning the transcriptional activity to these cell type categories. To make this latter point more concrete: while most photosynthesis in Arabidopsis takes place in mesophyll cells, other cell types also contain some low density of chloroplasts (Ishikawa et al., 2020). The increased transcriptional activity of photosynthetic processes in these other cell types at certain times of day may lead to an over-estimation of mesophyll transcriptional activity by CIBERSORTx, as photosynthesis related processes are used as markers for mesophyll transcriptional activity. We chose to use the term ‘cell-type specific transcriptional activity’ to highlight the fact that we are analysing the relative frequency of cell-type specific patterns of transcription. Despite the multiple different underlying processes that can lead to changes in cell-type specific transcriptional activity, as we define here, this concept is still very useful, because it provides us with a summary of the distinct transcriptional programmes that are occurring in samples that contain mixes of different cell types.

### Shifts in cell transcriptional activity over different time scales

We show that the transcriptional activity of several different cell types vary both diurnally and developmentally, as summarised in Fig 7. This suggests that waves of expression in bulk RNA-seq time series may represent waves of transcriptional activities of different cell types, rather than waves of regulatory activity– an assumption that is often held when inferring gene regulatory networks based on bulk RNA-seq data (Huynh-Thu and Geurts, 2018). Time series single-nucleus RNA-seq (Lee et al., 2023) and spatial transcriptomics (Giacomello, 2021) in plants promises to better distinguish temporal and spatial waves.

**Figure 7:**
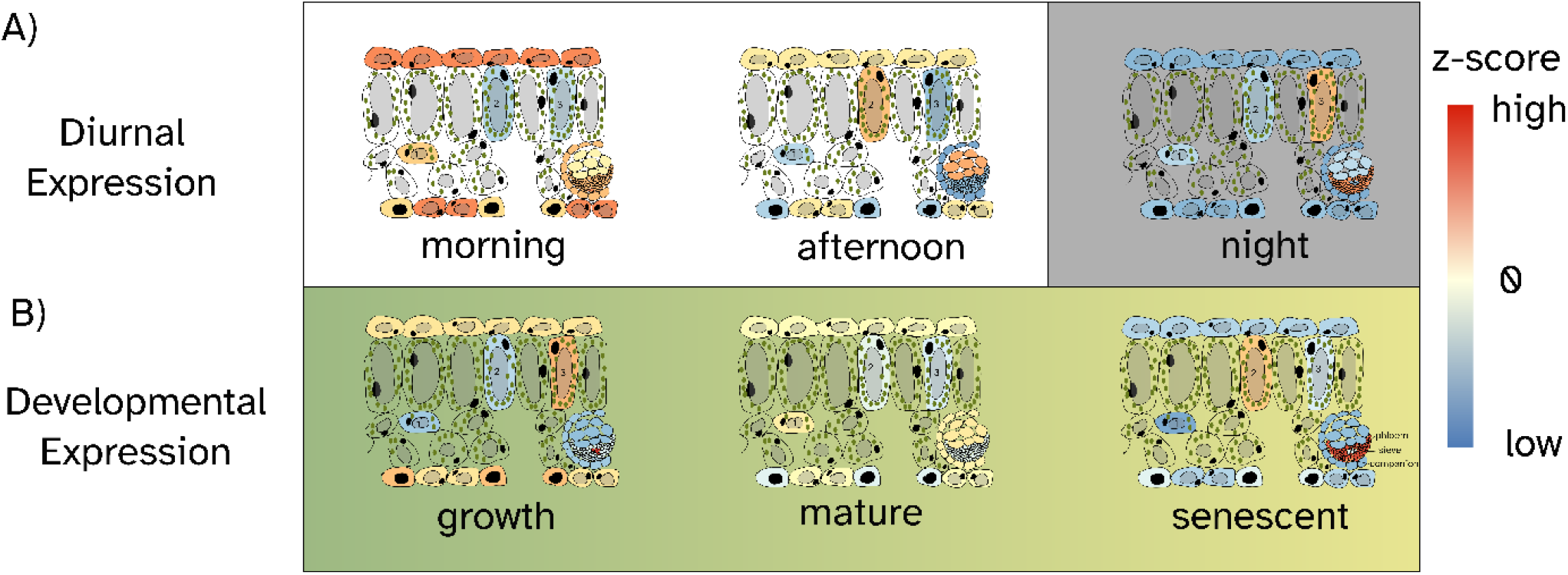
Graphical abstract showing the cell type-specific expression pattern across diurnal (A) and developmental (B) time series. The colours correspond the colour scale of the heatmaps in Figure 2.

### Novel patterns of mesophyll transcriptional states

Single-cell and single-nucleus RNA-seq has consistently identified sub-clustering of mesophyll cells, both in Arabidopsis and rice (Berrio et al., 2022; Xia et al., 2022; Kim et al., 2021; Procko et al., 2022; Wang et al., 2021). These studies have enabled the identification of cell type specific markers for different sub-groups of mesophyll; however, the main roles of these sub- clusters have not been fully characterised and the mesophyll groups do not have obvious physical differences. Our analysis augments Procko et al’s., (2022) analysis of the three mesophyll groups (as summarised in Table 1), by revealing the time in which each group is most likely transcriptionally active, both on a diurnal and developmental scale.

**Table 1.** Summary of properties of mesophyll cell types.

| Mesophyll | Period of development | Time of day | Light sensitive genes | Impact of bolting on relative expression |
| --- | --- | --- | --- | --- |
| 1 | Mature | Morning | LNK1, LNK3, STO, LHCB2, LHCA4. | Increase |
| 2 | Senescence | Afternoon to early evening |  | Increase |
| 3 | Growth | Night |  | Decrease |
| Stressed | Late senescence | 2-4 hours after dawn | DNAJ, SR45a, MBF1C | N/A |

It is important to remember that different parts of the same individual plant may perceive time in a different way, a form of intra-organismal heterochrony. Gould et al., (2018) found that there are waves of circadian expression that spread spatially from the meristem and root tips to the remainder of the plant, which could result in circadian asynchrony between cells in the leaf under free-running conditions. Asynchrony of the clock may also arise between cells in the leaf due to the entertainment of the clock in the morning through exogenous sugars (Haydon et al., 2013). There is also intra-organismal heterochrony in relation to biological age. Different leaves will be at different stages of development at the same chronological time (Efroni et al., 2008; Redmond et al., 2023). It may even be possible for different parts of the same leaf to have different biological ages at the same chronological age, due to localised stresses. The individual transcriptional profiles sampled using single cell or single nucleus RNA-seq methods may not only represent heterogeneous cell types but also heterogeneous biological times. By combining single cell RNA-seq data with high temporal resolution bulk RNA-seq, we may be able to begin to dissect this kind of temporal heterogeneity too.

### CIBERSORTx enables greater exploitation of RNA-seq datasets

While single cell RNA-seq is becoming increasingly common in plants, it is still too expensive and cumbersome to perform these experiments over high resolution time series, under a wide range of environmental conditions, and under a wide range of genotypes. In addition, there are tens of thousands of existing bulk RNA-seq datasets available that could vary in their cell type proportion (Zhang et al., 2020). Here we show that plant scRNA-seq datasets can be used to train a CIBERSORTx model that can be used to algorithmically dissect the bulk RNA- seq samples by their cell types. This will enable Arabidopsis researchers who do not have the capacity to do scRNA-seq the ability to exploit this data for enhancing their research. Moreover, AraLeTA is a useful platform for utilising existing bulk RNA-seq time series data, single cell RNA-seq data, and transcription factor binding data to isolate spatial and temporal regulatory processes of interest. Together, CIBERSORTx and AraLeTA provide us with an atlas of leaf expression in Arabidopsis over cell type and over time.

## Methods

### Running CIBERTSORTx

Seurat was used to visualise and process the single cell RNA-seq data (Hao et al., 2021). In the matrix provided to CIBERSORTx (Newman et al., 2019) to generate the signature matrix, genes that were very lowly or very highly expressed were filtered out, with ln(total read count) between 4 and 10. For the purposes of testing CIBERSORTx on the simulated bulk-RNA-seq data, a random sample of 75% of the cells were used to generate the signature matrix. For the rest of the paper, all cells were used to generate the signature matrix. The cluster designations used in Procko et al. (2022) were used as the phenotype classes. In all cases, we disabled quantile normalisation (which is recommended for RNA-seq data) and we used 100 runs for the permutation tests. For imputing the gene expressions in each sample in the bolted plants, we used only the 9 most abundant cell types and “other” and we focussed only on genes that were identified as significantly up or down regulated in bolted/not bolted plants, according to Redmond et al., (2023).

### Generating simulated bulk RNA-seq samples

To simulate bulk RNA-seq samples composed on a single cell type, the remaining cells from Procko et al., (2022) that were not used to generate the model were randomly partitioned into two equal groups for each cell type and summed together, forming replicate 1 and 2 for each cell type. We also simulated bulk RNA-seq samples from the single cell microdissected samples from Xia et al., (2022). In this case, we summed over the single cell RNA-seq samples from each cell type associated with a specific leaf.

To simulate the mixed bulk RNA-seq samples, we wanted to have somewhat realistic cell type proportions, but to also have a wide variation in cell types between samples. For each cell type, we first calculated the proportion of cells of that type in the single cell RNA-seq dataset. Then, per cell type, we simulated the proportion of cells within the bulk RNA-seq sample according to a normal distribution, with a mean and standard deviation equal to the cell type proportion within the single cell RNA-seq data set. To calculate the final number of cells per cell type, we rounded up any negative values to 0, and multiplied this by the average number of cells we wanted per sample (600). We randomly selected the specified number of cells from each cell type and then calculated the sum of transcript counts per gene for the associated set of cells, and considered that to be our simulated bulk-RNA-seq sample.

### Bioinformatics analysis

All clustering was performed using the default hierarchical clustering parameters of the pheatmap package (Kolde, 2019). Networks were visualised using Gephi (Bastian, et al., 2009) and iGraph (Csárdi et al., 2023). PAFway (Mahjoub and Ezer, 2020) was used to generate a graph of the related functional annotations of the basis of the text analysis of the Gene Ontology Slim annotations provided by Berardini et al., (2015) and the topology of the DAP-seq gene network (O’Malley et al., 2016). AraLeTA was developed as a Shiny App (Chang et al., 2023). The PAFway networks were visualised using Gephi (Bastian et al., 2009a). All code for generating the figures is available on https://github.com/stressedplants/AraletaScripting/. All code for the Shiny app is available on: https://github.com/stressedplants/AraLeTA/.

## Acknowledgements

We would like to acknowledge the following funding sources: Royal Society (RGS\R2\212345, Ezer), BBSRC IAA (BB/S506795/1, Ezer), BBSRC Responsive Mode (BB/V006665/1, Ezer, Davis), the BBSRC White Rose DTP (BB/T007222/1, Vong, Redmond) and GenerationResearch (Claydon).

## Author contributions

Ezer designed the research. All authors performed research. Ezer and Vong contributed new analytic/computational tools. Ezer, Vong, McCarthy and Claydon analysed data. All authors wrote the paper.

