## Supplemental Materials for "An Arabidopsis leaf expression atlas across diurnal and developmental scales"

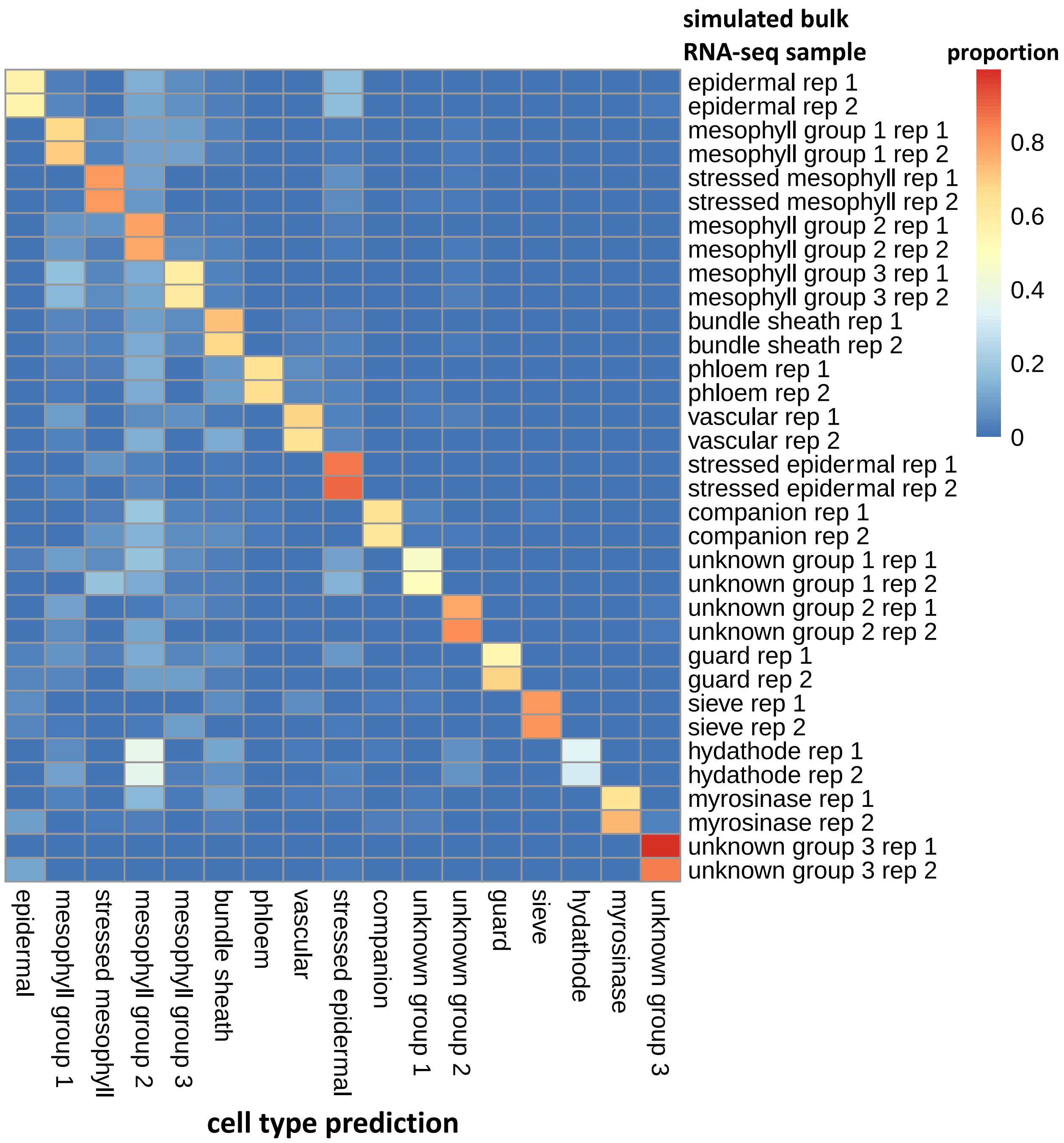


**Figure S1: Ability of CIBERSORTx to detect 'mock' bulk RNA-seq samples, consisting of a single cell type.** (A) The cluster designations from Procko et al. are used throughout (2022). CIBERSORTx was trained on 75% of the cells from each of these clusters. The remaining cells were used to develop simulated bulk RNA-seq samples. For each cluster, we generated two simulated bulk RNA-seq samples of pure tissue, by summing the expression of 50% of the cells from that cluster that were not used for training. (B) The heatmap shows the proportion of each cell type (columns) that was predicted for each simulated bulk RNA-seq sample (rows). The simulated RNA-seq samples are labelled by the cell type that was used to generate the simulated bulk RNA-seq sample.

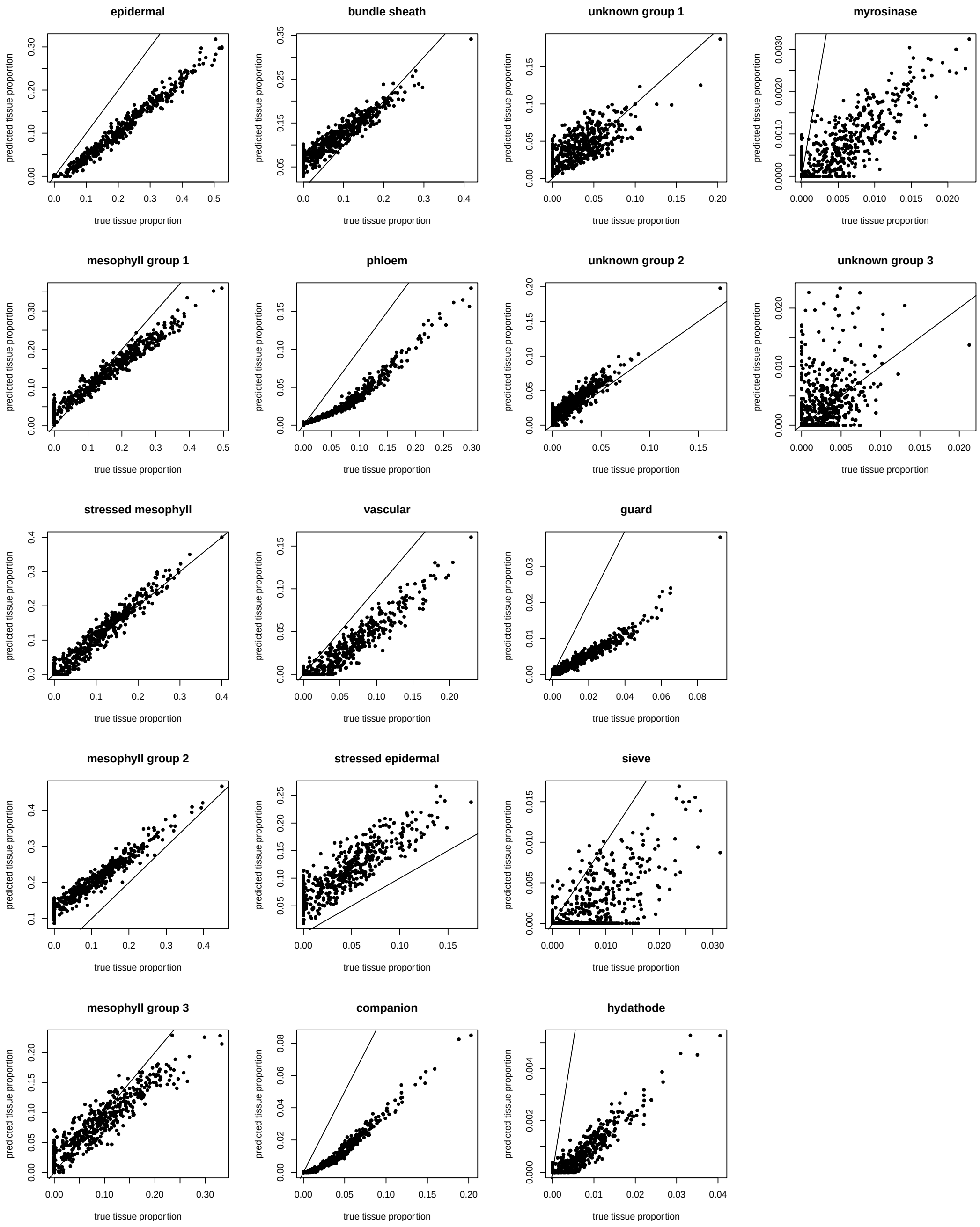


**Figure S2: Ability of CIBERSORTx to predict relative composition of 'mock' bulk RNA-seq samples with mixed tissues.** 75% of the cells per cluster were used to train a CIBERSORTx model, while the remaining 25% of cells were repeatedly sub-sampled to generate 500 simulated bulk RNA-seq samples with mixed cell types (see Methods for sampling strategy). These graphs compare the true proportion of each cell type in the simulated bulk RNA-seq sample with the proportion predicted by CIBERSORTx. The line on each sub-plot has a slope of 1 and a y-intercept of 0.

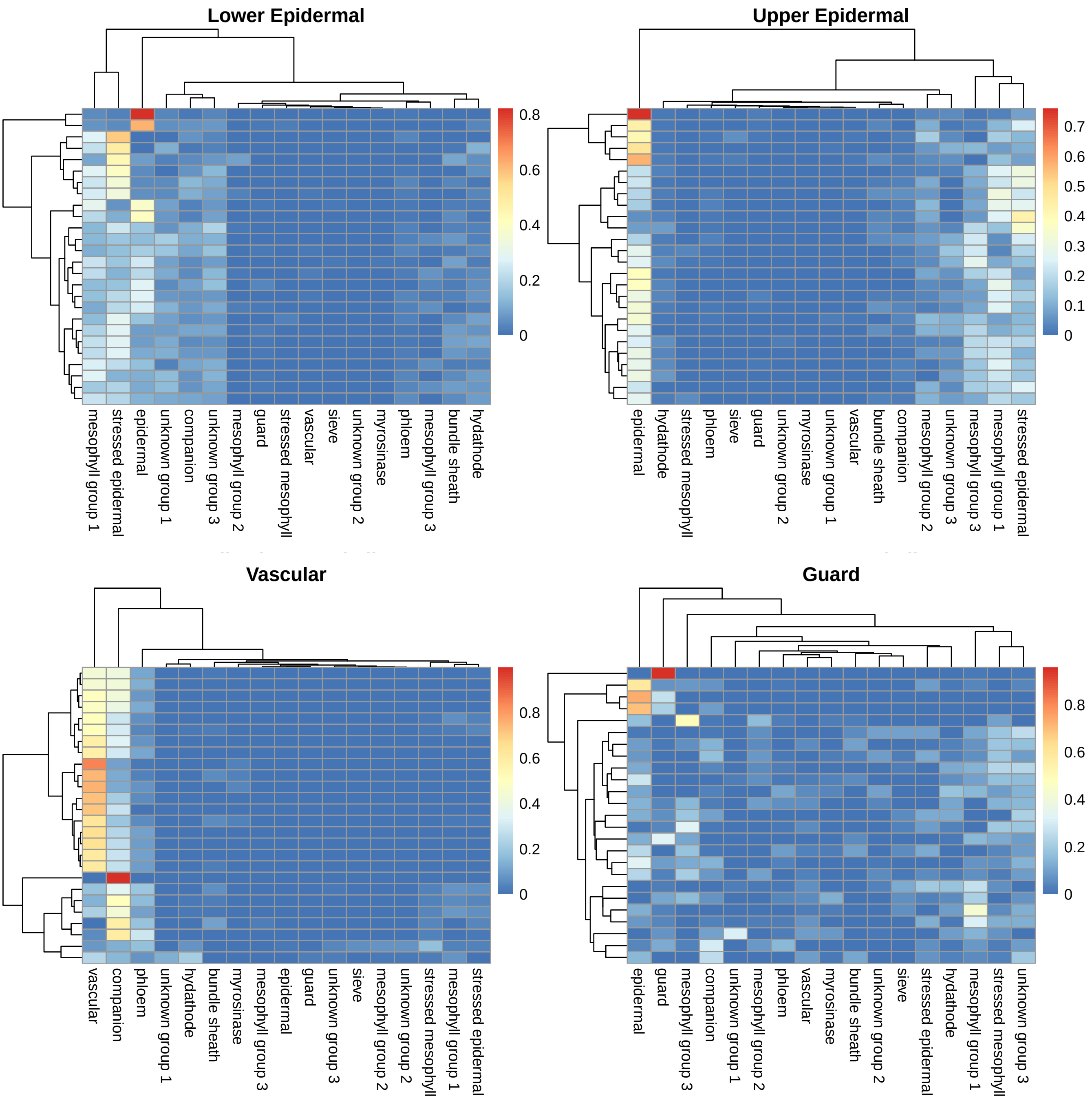


**Figure S3: Assignment of cell type to microdissected tissues.** Here we estimate the proportion of cell types of microdissected by Xia et al., (2022) on the basis of the scRNA-seq data from Procko et al. (2022), using CIBERSORTx. This heatmap clusters the predicted cell type proportions in microdissected leaf samples, with each row representing a leaf and each column representing a different predicted cell type.

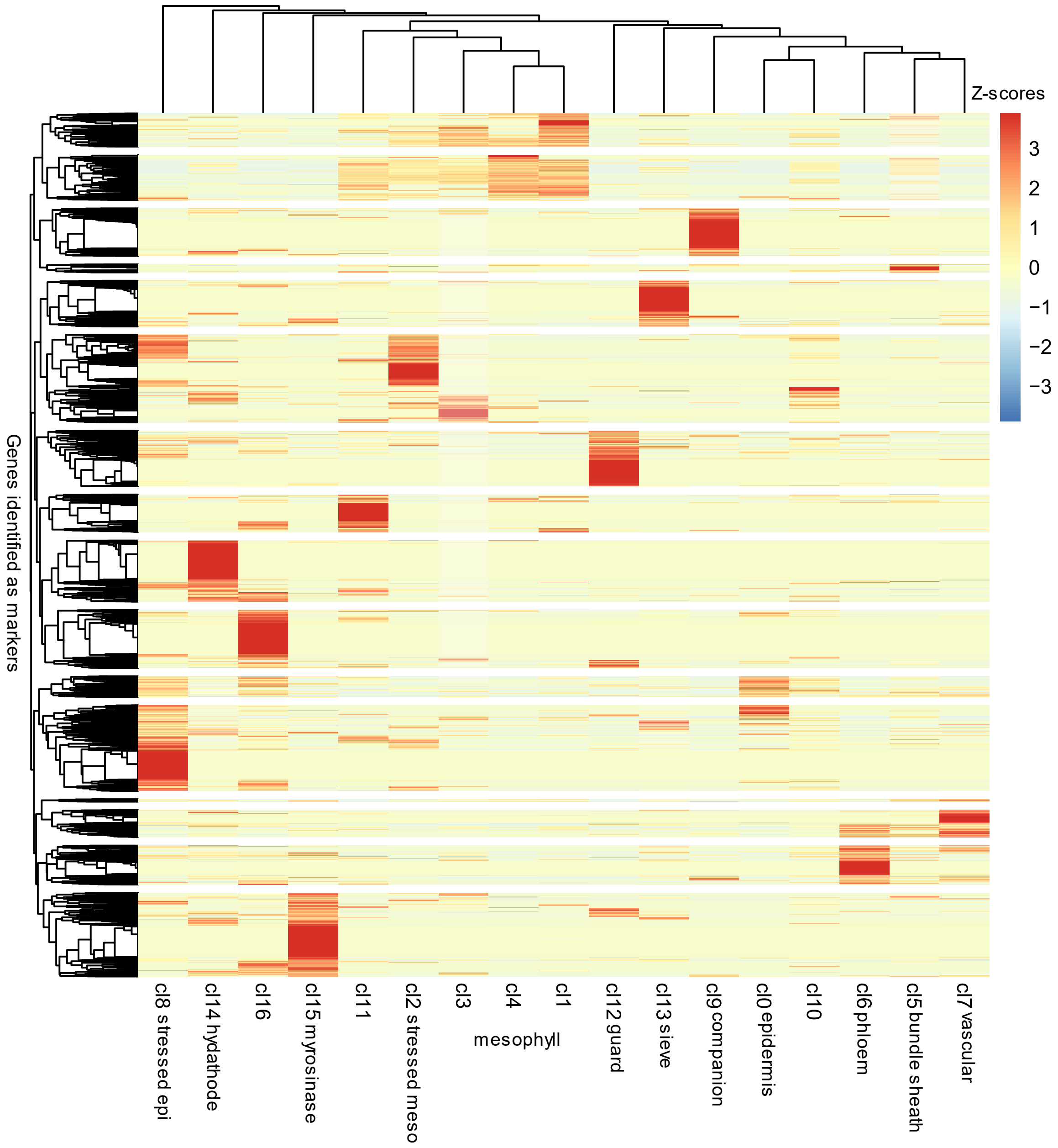


**Figure S4: Analysis of signature matrix.** This is a clustering of the signature matrix that is used by CIBERSORTx to dissect the proportions of cell types, with each row normalised. Please note that most cell types appear to have a unique signature, but mesophyll cluster 4 is distinguished by a lack of expression of certain mesophyll genes and that there is only a small set of genes that distinguish unknown cl10 and bundle sheath cells.

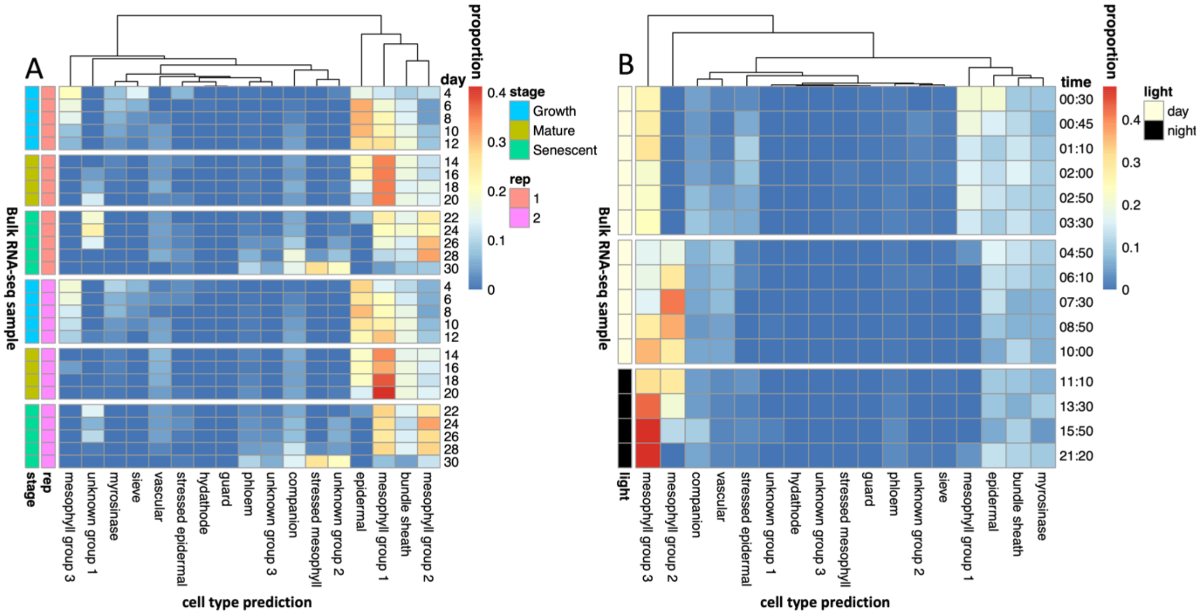


**Fig S5: Unscaled heatmaps from Fig 2**. Here we show the same heatmaps as Fig 2, but showing the actual proportions of cell types predicted, rather than the z-scores. Recall from Fig 1 that CIBERSORTx is not very reliable at predicting the relative expression of cell types within a sample, but is good at predicting this between samples, which is why the z-scores are preferable to use.

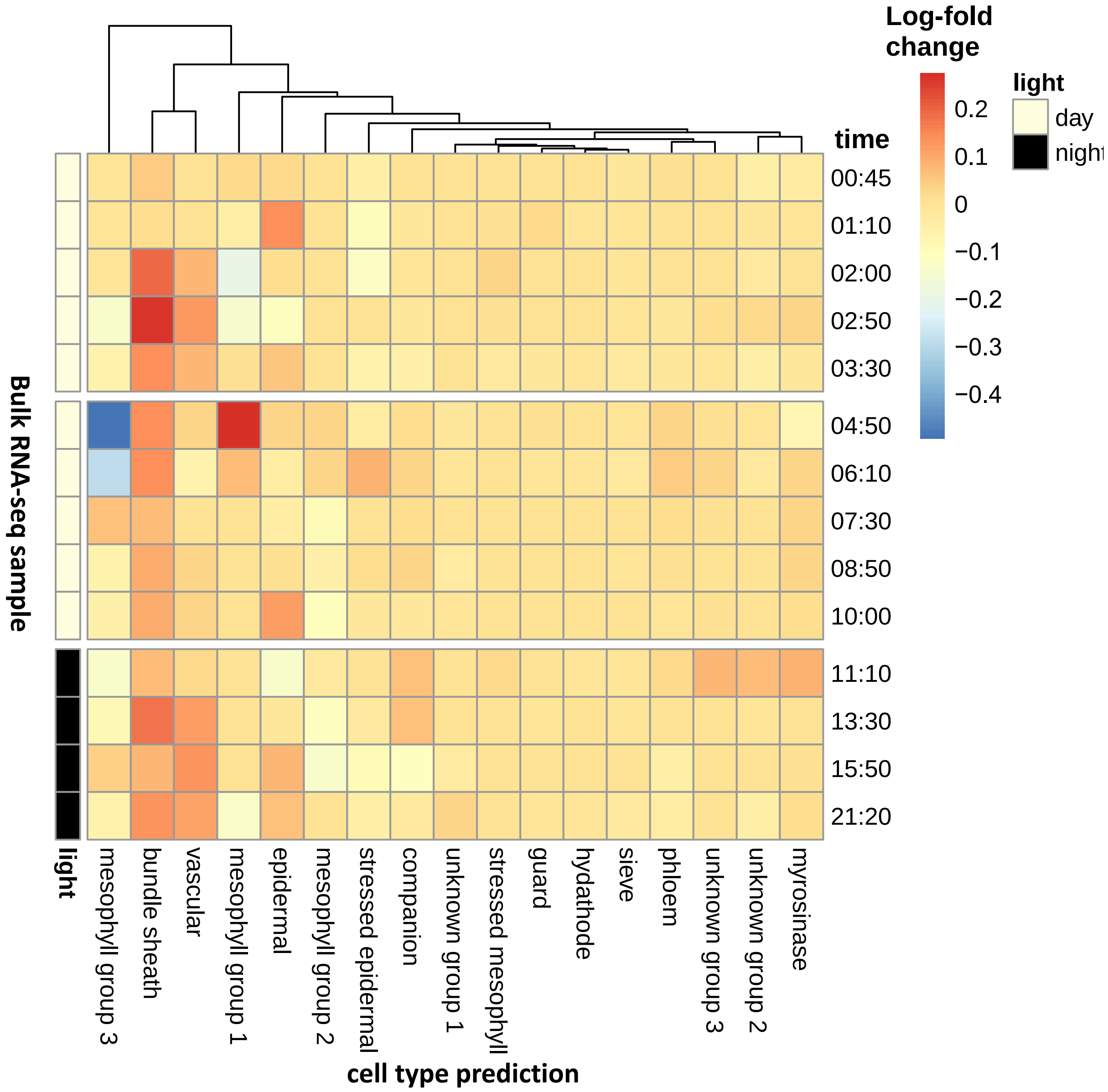


**Figure S6: Impact of MeJA on cell type proportions.** Log-fold change in predicted proportion of cell types in cells exposed to MeJA vs controls that were mock treated at the same time point.

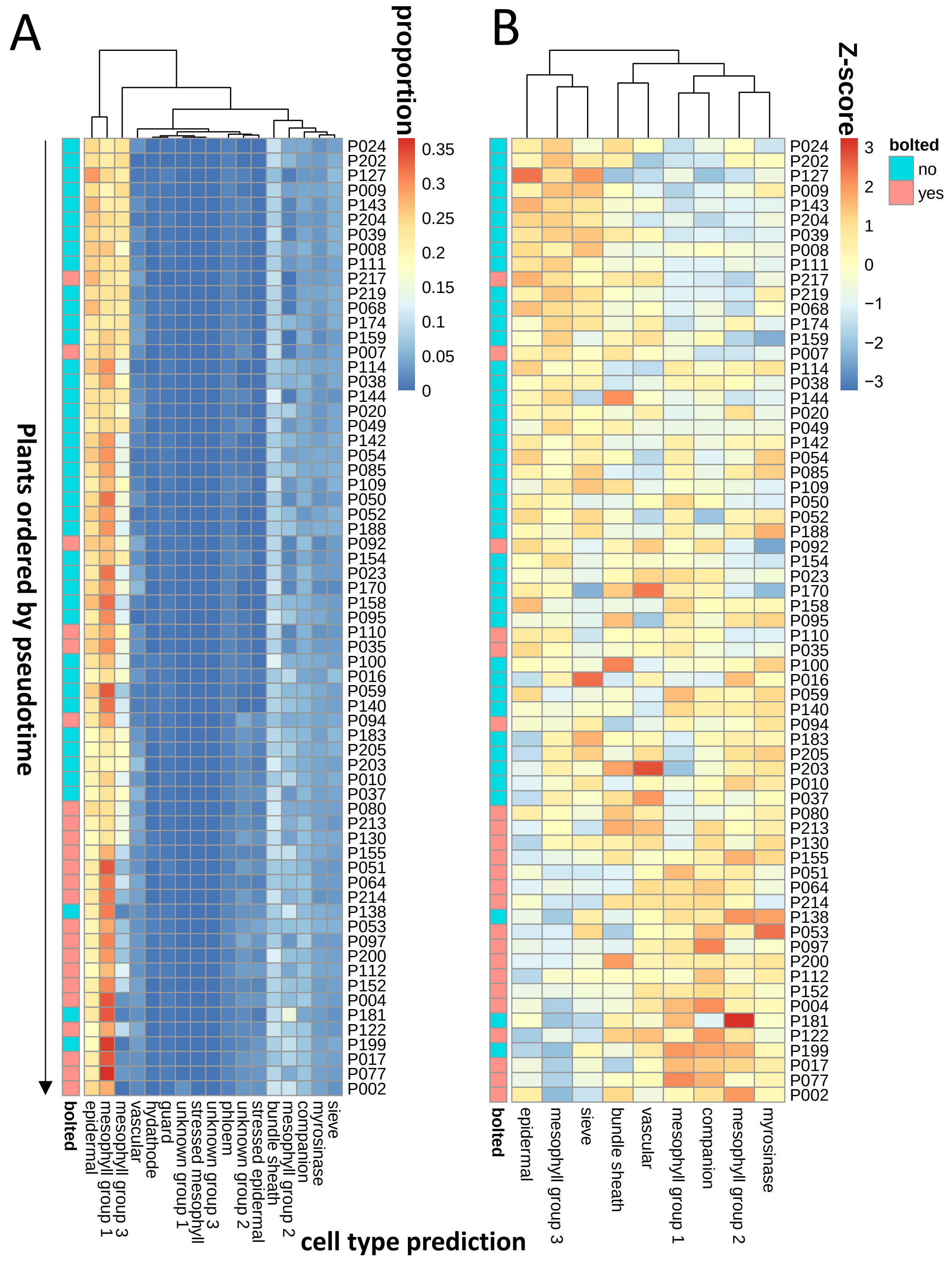


**Figure S7: Unscaled heatmaps from Fig 4**. Here we show the same heatmaps as Fig 4, but showing the actual proportions of cell types predicted, rather than the z-scores. Recall from Fig 1 that CIBERSORTx is not very reliable at predicting the relative expression of cell types within a sample, but is good at predicting this between samples, which is why the z-scores are preferable to use.

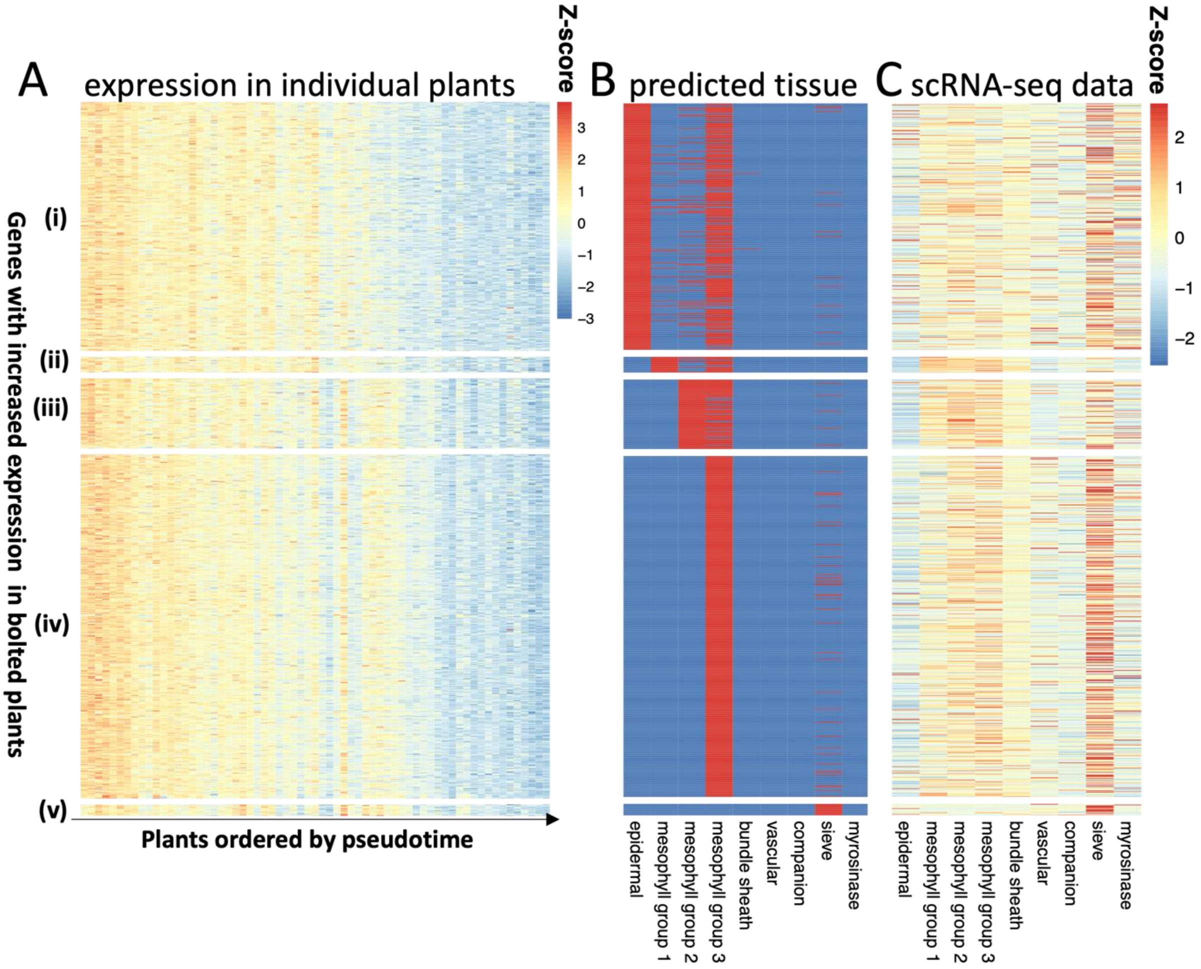


**Figure S8: Cell-type specific gene expression of genes that have low expression in leaves of bolted plants.** Using the high-resolution imputing function in CIBERSORTx, we predicted cell-type specific expression of genes that were differentially expressed in bolted/unbolted plants. Here we show the results for the genes that have lower expression in bolted plants. (A) shows the z-score of the expression of these genes in the bulk RNA-seq experiment (Redmond et al., 2023), grouped by the cell type in which they were predicted to be expressed and ordered by pseudotime. (B) shows the cell type assignment, with red indicating that a gene is expected to be found in that cell type. (C) shows the z-score of the mean expression of these genes in the scRNA-seq data set (Procko et al., 2022).

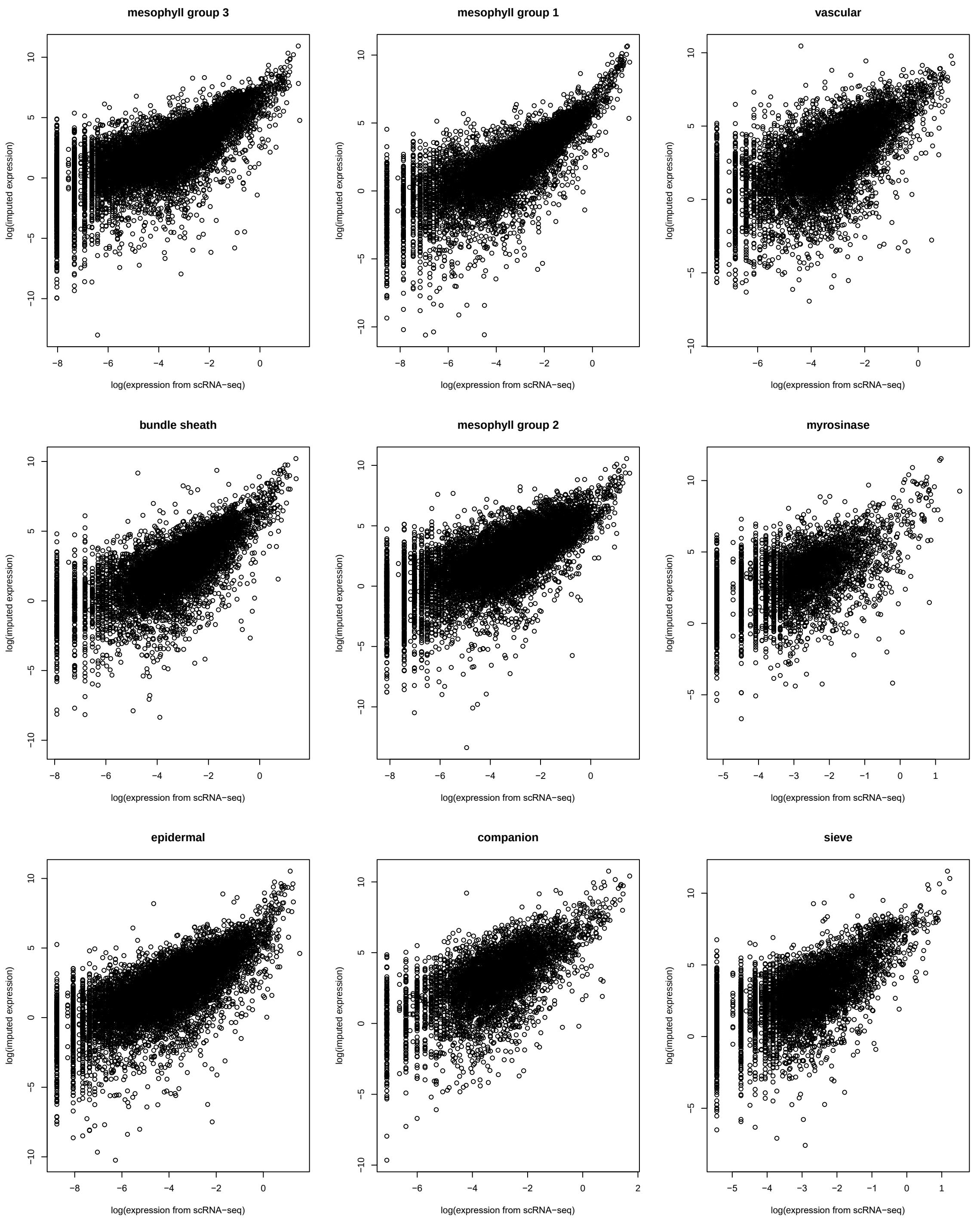


**Figure S9: Comparison of predicted gene expression values and imputed values**. For each cell type, we compared the gene expression values from Procko et al., (2022) and our predicted values. Note that CIBERSORTx only uses a small set of genes for predicting cell type proportions, so it does not have access to information about the cell type specific expression of most genes during the imputing process.

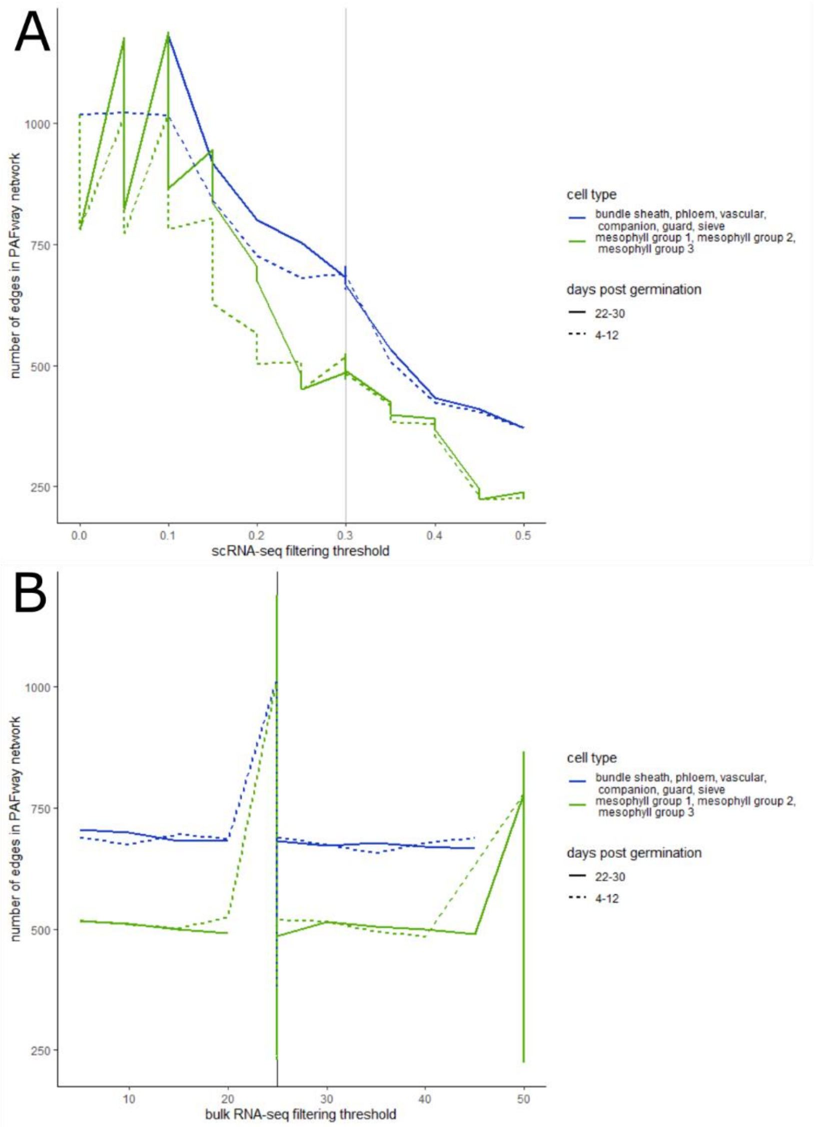


**Figure S10: Identification of scRNA and bulk RNA thresholding values**. (A) For each of the four conditions under investigation, the scRNA-seq thresholds were adjusted between 0 and 0.5, while the bulk RNA-seq threshold was set to 25, and the number of significant edges ($p<0.05$) in the PAFway network were calculated. In all of the cases, there was a local optimum or an inflection point at 0.3, so this parameter value was selected. (B) Then, the bulk RNA-seq thresholds were adjusted between 0 and 50 and local optima were chosen.

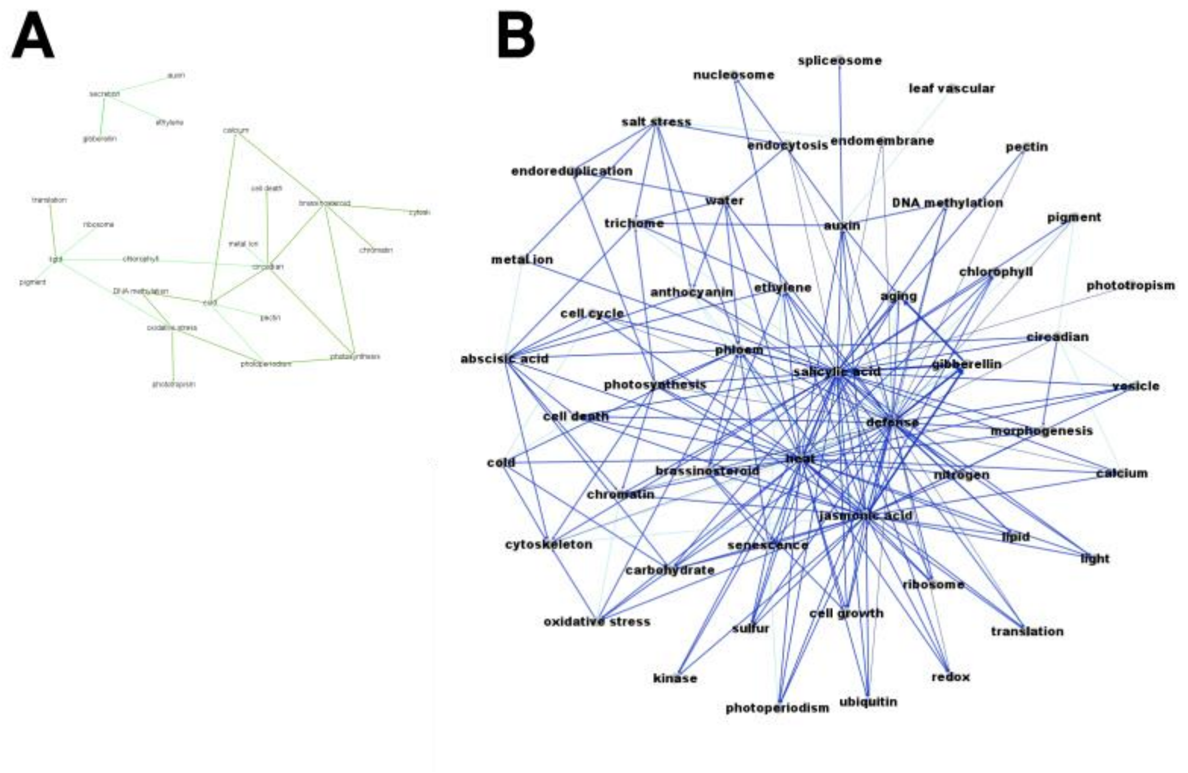


**Figure S11: Comparison of predicted gene expression values and imputed values**. Figure 6B and 6C show the networks for old and young plants, colour-coded by whether they appear in mesophyll cells, vascular cells or both. Here we show the complementary networks, split by cell type. (A) The PAFway network for mesophyll cells are shown. The colours of the edges correspond to the age, as shown in the Venn Diagram in Figure 6A. (B) The PAFway network for vascular cells are shown.

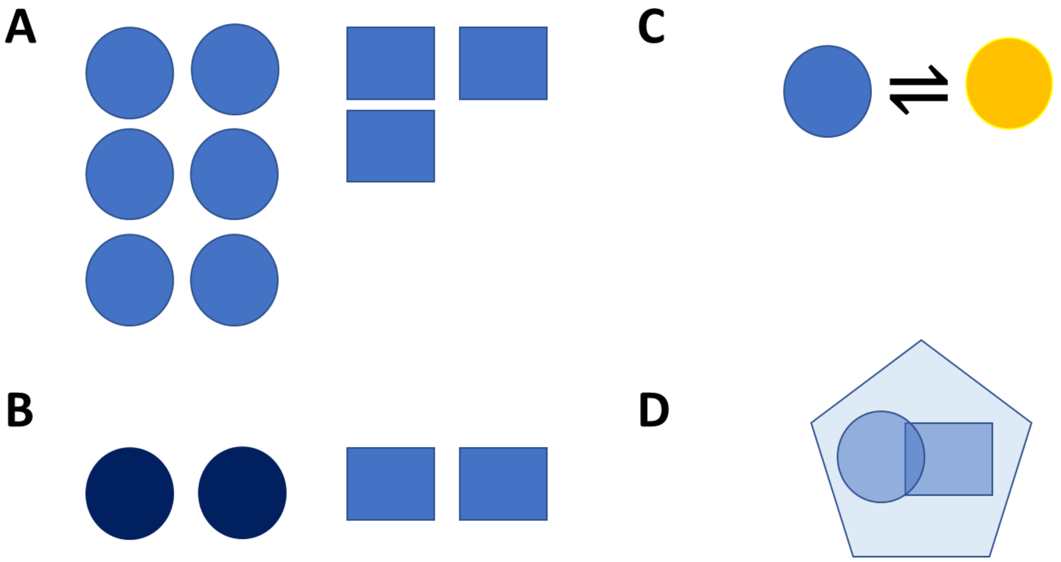


**Figure S12: Factors influencing transcriptional activity**. CIBERSORTx infers relative cell-specific transcriptional activity. There are several different mechanisms that can generate discrepancies between cell-specific transcriptional activity, including: (A) different proportions of cell types between samples, (B) different transcriptional rates within the cell types sampled (C) cell types changing state within sample (D) unique cell type with transcriptional activity resembling combinations of other cell types.
